# The influence of higher-order epistasis on biological fitness landscape topography

**DOI:** 10.1101/164798

**Authors:** Daniel M. Weinreich, Yinghong Lan, Jacob Jaffe, Robert B. Heckendorn

## Abstract

The effect of a mutation on the organism often depends on what other mutations are already present in its genome. Geneticists refer to such mutational interactions as epistasis. Pairwise epistatic effects have been recognized for over a century, and their evolutionary implications have received theoretical attention for nearly as long. However, pairwise epistatic interactions themselves can vary with genomic background. This is called higher-order epistasis, and its consequences for evolution are much less well understood. Here, we assess the influence that higher-order epistasis has on the topography of 16 published, biological fitness landscapes. We find that on average, their effects on fitness landscape declines with order, and suggest that notable exceptions to this trend may deserve experimental scrutiny. We explore whether natural selection may have contributed to this finding, and conclude by highlight opportunities for further work dissecting the influence that epistasis of all orders has on the efficiency of natural selection.

## 1 Introduction

One of the more evocative pictures of evolution is that of a population climbing the fitness landscape [41, 48]. This image was originally proposed by Sewall Wright [81] to build intuition into his [80] and R.A. Fisher’ s [22] technical treatment of Darwin’ s theory of natural selection in finite populations under Mendelian genetics [55]. The topography of the fitness landscape represents the strength and direction of natural selection as local gradients that influence the direction and speed with which populations evolve.

While several distinct framings of the fitness landscape have been suggested [55], here we employ the projection of genotypic fitness over Maynard Smith’ s sequence space [40]. Sequence space is a discrete, high-dimensional space in which genotypes differing by exactly one point mutation are spatially adjacent. Thus, proximity on the fitness landscape corresponds to mutational accessibility, and selection will try to drive populations along the locally steepest mutational trajectory. (See [75] for several processes not readily captured by this construction.)

The most obviously interesting topographic feature of the fitness landscape is the number of maxima, a point already recognized by Wright [81]. Two (or more) maximum can constrain natural selection’ s ability to discover highest-fitness solutions, since populations may be required to transit lower-fitness valleys on the landscape en route. (Though see [29, 72] for the population genetics of that process, sometimes called stochastic tunneling [15, 29].)

### 1.1 Epistasis and fitness landscape topography

Epistasis is the geneticist’ s term for interactions among mutational effects on the organism [50]. For example, genetically disabling two genes whose products act in the same linear biochemical pathway can have a much more modest effect than the sum of the effects of disabling either gene in isolation. Conversely, disabling two functionally redundant genes can have a much more substantial effect than expected. (Indeed, such observations have taught us quite a bit about the organization of biochemical pathways, e.g., [2].)

Epistatic interactions between mutations can occur for any organismal trait, including fitness. Importantly, epistasis for fitness has an intimate connection to the topography of the fitness landscape, a fact also already appreciated by Wright [81]. For example, multiple peaks require the presence of mutations that are only conditionally beneficial (called sign epistasis [53, 75]). More generally, an isomorphism exists between fitness landscapes defined by mutations at some *L* positions in the genome and the suite of epistatic interactions possible among them. This follows because, while any particular mutation can appear on 2^*L*-1^ different genetic backgrounds (assuming two alternative genetic states at each position), each such mutation-by-background pair corresponds to a distinct adjacency in sequence space. Consequently, arbitrary differences in the fitness effect of a mutation across genetic backgrounds (i.e., epistasis of all order) can generically be represented on the fitness landscape [75].

### 1.2 Higher order epistasis and natural selection

Widespread epistasis between pairs of mutations has been recognized in nature for over 100 years [50, 74], and the corresponding theory is fairly advanced (e.g., [5,79]). However, interactions can themselves vary with genetic background, called higher-order epistasis [13, 74]. And while it is now becoming clear that higher-order interactions are commonplace in nature [36, 46, 68, 74], their influence on natural selection is less well understood (though see [59]). Clearly, epistatic interactions of all order can render natural selection less efficient (e.g., [75, 79]). It is thus reasonable to suppose that higher-order epistasis might be particularly likely to confound selection’ s ability to efficiently improve fitness. To test this idea, we relied on the fact that empirically observed fitness landscapes are sampled by virtue of their occupancy by natural populations, which are themselves the products of natural selection. This motivated our specific hypothesis: that epistatic influence on the topography of naturally occurring fitness landscapes should decline with epistatic order. We tested this prediction using 16 published biological fitness landscapes.

## 2 Methods

### 2.1 The order of epistatic interactions

Any set of *L* biallelic loci defines 2^*L*^ genotypes, each with 2^*L*^ potentially independent fitness values. Simultaneously, there are 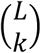 distinct subsets of *k* mutations that in principle can also independently contribute to a genotype’ s fitness. In total, there are thus 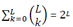 subsets of mutations (i.e., the power set of *L* mutations). This counting reflects the isomorphism between any fitness landscape and its corresponding suite of epistatic terms [74].

We designate interactions among any subset of *k* mutations as *k*^th^-order epistasis. Note that here first-order “ epistasis” is degenerate in the sense that it represents the fitness effects of each of the *L* mutations in isolation. And our zeroth-order “ epistatic” term is the benchmark, fitness landscape-wide mean, relative to which the effect of each subset of mutations is computed.

### 2.2 The Fourier-Walsh transformation

Following earlier work [26, 45, 65, 71, 74] we employ the Fourier-Walsh transformation (Fig. 1a) to convert between fitness landscapes and their corresponding epistatic terms. This is a linear transformation written

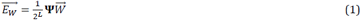

**Fig. 1.**
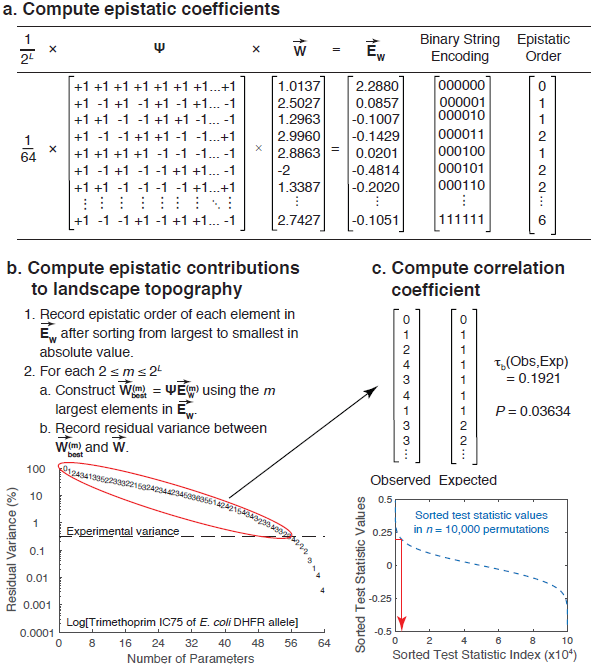
Analytic pipeline illustrated with data from Palmer et al., 2015 [49]. **a.** For each dataset, fitness data (or a suitable proxy) were first converted to the corresponding epistatic terms using the Fourier-Walsh transformation (Eq. 1). **b.** Predicted fitness under a succession of models using only the *m* largest epistatic terms in absolute value were compared with the published data. For given value of *m*, these models provably have the greatest explanatory power (smallest residual variance) of any model with exactly *m* parameters (Appendix). The symbols plotted represent the epistatic order of each successive parameter added to the model. **c.** Rank correlation coefficient between the sequence of epistatic orders observed and those of our naïve expectation (Eq. 2) were computed.Statistical significance was assessed by a permutation test that asked whether the observed sequence of epistatic orders was significantly different than random.

Here 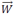 is the vector of all 2^*L*^ fitness values arranged in the canonical order defined by ascending *L*-bit binary numbers encoding the corresponding genotype with respect to the presence or absence of each mutation (e.g., [37]). (*W* is the traditional population genetics symbol for fitness.) Ψ is the Hadamard matrix, the unique, symmetric 2^*L*^ × 2^*L*^ matrix whose entries are either +1 or -1 and whose rows (and columns) are mutually orthogonal. Finally, 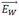is the resulting vector of 2^*L*^ epistatic terms arranged in the canonical order defined by ascending *L*-bit binary numbers whose 1’ s indicate the corresponding subset of interacting loci. Fig. 1a illustrates this transformation. (See [54] for the relationship between this and other formalisms for computing epistatic terms.)

The orthogonality and symmetry of Ψ means that Ψ^T^ · Ψ = Ψ^2^ = 2^*L*^**I**, where **I** is the identity matrix. This means that, just as Eqn. 1 converts any landscape into its epistatic terms, so too can any vector of epistatic terms E be converted into its corresponding fitness landscape as 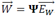 WEtake advantage of this fact next.

### 2.3 Subsetting approximations of a fitness landscape

Given fitness function 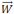 we now introduce subsetting approximations 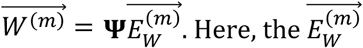 are constructed so as to have 0 ≤ *m* ≤ 2^*L*^ of the components in 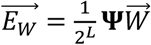 (Eqn. 1) and the remaining 2^*L*^ *– m* components set to zero. There are thus 2^2L^ subsetting approximations for any fitness function 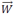 (corresponding to the power set of the 2^*L*^ epistatic terms in 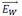. As a consequence of the orthogonality of the Fourier-Walsh transformation, the sum of squares distance between fitness function 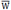and subsetting approximation 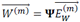is minimized for given *m* if and only if 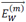uses the *m* largest components in absolute value of 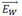 (Appendix 1). We denote these 0 ≤ *m* ≤ 2^*L*^ best subsetting approximations 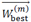. (Subsetting approximations defined by interaction order rather than absolute magnitude of epistatic terms were recently employed elsewhere [59].)

### 2.4 Quantifying the influence of epistatic terms on empirical fitness landscape topography

To examine the influence of epistasis on fitness landscape topography as a function of epistatic order, we first used Eqn. 1 to compute 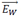 for each 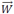gleaned from the literature (§2.7). For each 1 ≤ *m* ≤ 2^*L*^, we then iteratively constructed each 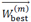 Finally, for each *m* we recorded the residual variance between 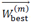 and 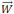(minimized by this subsetting approximation; §2.3), together with the epistatic order of the *m*^th^-largest component of 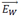. Fig. 1b illustrates this process.

### 2.5 Statistics

Our hypothesis is that the influence of an epistatic term on the fitness landscape should decline with epistatic order. Put another way, we expected that after sorting the elements of 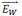 (Eqn. 1) by their absolute magnitudes, the associated epistatic orders should be represented by a vector of 2^*L*^ integers that reads:

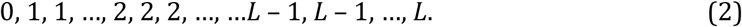

Specifically, this vector consists first of one zero followed by *L* ones, 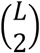 twos and in general 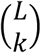*k*’ s for all 1 ≤ *k* ≤ 2^*L*^.

We tested this hypothesis for each dataset by first computing Kendall’ s τb correlation coefficient [31] between this expectation and the epistatic orders observed among the elements in 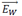 sorted by absolute magnitude. τb is one (negative one) when the observed epistatic orders are perfectly correlated(anticorrelated) with expectation, and zero when they are uncorrelated. Note that Kendall’ s τb statistic is appropriate because it accommodates ties. For studies that also reported experimental variance, we computed the correlation coefficient after discarding the epistatic orders of all *j* elements in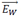 that reduced residual variance by less than experimental variance (Fig. 1b, Table 1) as well as the last *j* epistatic order values in our expectation (Eqn. 2).

**Table 1.**
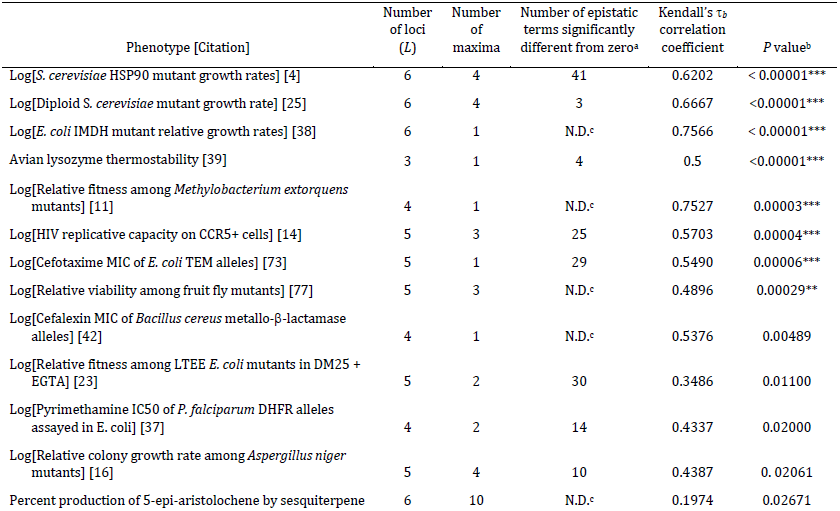
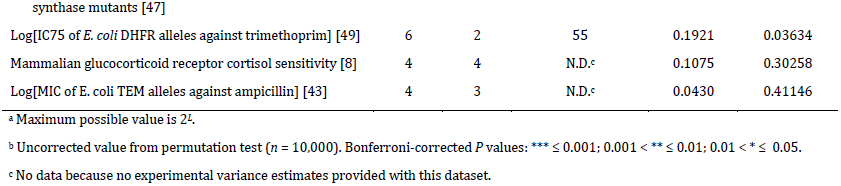
Analyses of published combinatorially complete fitness landscapes

For each dataset, we then used a permutation test to test the null hypothesis that the corresponding correlation coefficient is zero. Specifically, each dataset is characterized by some number of epistatic terms: 2^*L*^ in cases where no experimental variance estimate is provided, or 2^*L*^ – *j* in cases where we were able to identify non-significant epistatic components (see previous paragraph and Table 1). For each of *n*= 10,000 replicates, we computed the rank correlation coefficient between two random permutations of this number (2^*L*^ or 2^*L*^ – *j*) of the epistatic order values drawn from Eqn. 2 for given *L*. We then sorted correlation coefficients, and the uncorrected *P* value reported for each dataset (Table 1) was taken as the fraction of permutations in which a correlation coefficient greater than or equal to the empirical value was observed (Fig. 1c). Thus, ours is a one-tailed test of the hypothesis that no positive correlation is present.

We used the Bonferroni-Holm method [28] to correct for multiple tests. In addition, under the null hypothesis that epistatic orders are uncorrelated with our naïve expectation, the distribution of *P* values observed across datasets should be uniformly distributed. We tested this hypothesis with a *G*-test after binning counts of empirically observed *P* values. We assessed statistical significance relative to the χ^2^ distribution [61].

### 2.6 Computing Kendall’ s correlation coefficient with aggregated higher-order epistatic terms

We also computed Kendall’ s correlation coefficient for the sequence of epistatic orders previously reported (Fig. 2 in [49]), in which third- and higher-order epistatic terms were not distinguished. To do this we coded all observed epistatic terms in this aggregate group as third-order. Those authors found that the residual variance in a model containing just 70 epistatic terms was roughly equal to the experimental variance. Thus, the analog to Eqn. 2 now contains one zero, followed by six ones,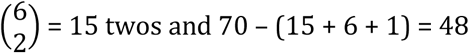 threes. Statistical significance was again assessed using a permutation test (*n* = 10,000) using this modified expectation in place of Eqn. 2.

**Fig. 2.**
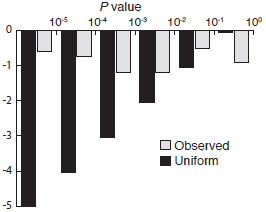
Distribution of uncorrected *P* values among 16 empirical datasets. Eight of 16 *P* values are statistically significant after correcting for multiple tests, and distribution of *P* values is sharply skewed toward small values.

### 2.7 Empirical datasets

To compute all 2^*L*^ epistatic terms in a fitness landscape defined over *L* biallelic loci requires data on the fitness values (or suitable proxy) for each of the corresponding 2^*L*^ genotypes. We previously designated such datasets combinatorially complete [74], and the datasets analyzed here are shown in Table 1. Several datasets [4, 38, 47, 49] had a few loci with cardinality greater than two. In these cases, we examined one “ slice” through the landscape defined by randomly choosing just two alleles at those loci.

Several studies examined multiple phenotypes for a single set of mutations, and follow-up studies sometimes presented additional phenotypes for a previously described set of mutations. Those cases are enumerated in Table 2; for each set of mutations we randomly sampled just one phenotype. Table 2 also lists all combinatorially complete datasets we know that are defined over loci with cardinality greater than two. These were excluded here because the Fourier-Walsh framework doesn’ t readily generalize to higher cardinalities.

**Table 2.**
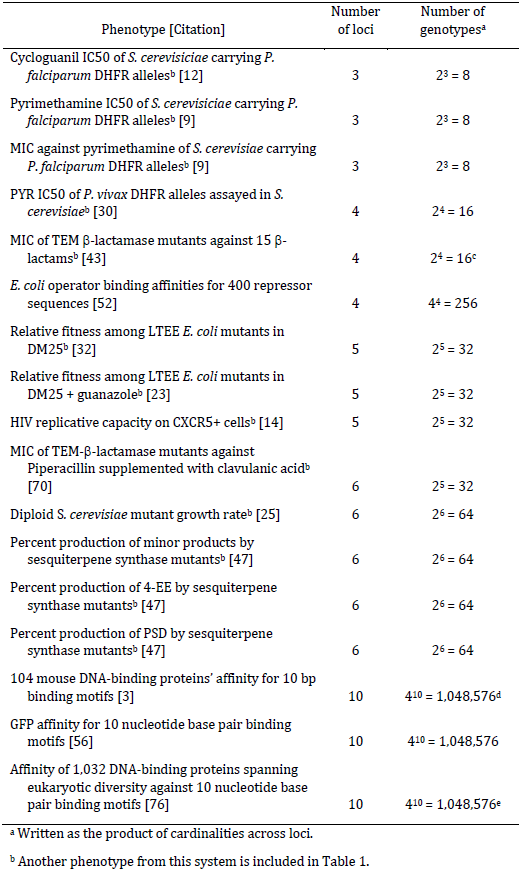
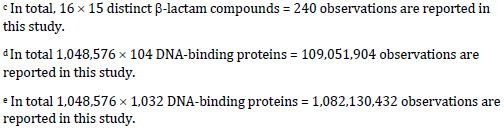
Published combinatorially complete fitness landscapes not examined here

| Phenotype [Citation] | Number of loci | Number of genotypes<sup>a</sup> |
| --- | --- | --- |
| Cycloguanil IC50 of *S. cerevisiae* carrying *P. falciparum* DHFR alleles<sup>b</sup> [12] | 3 | $2^3 = 8$ |
| Pyrimethamine IC50 of *S. cerevisiae* carrying *P. falciparum* DHFR alleles<sup>b</sup> [9] | 3 | $2^3 = 8$ |
| MIC against pyrimethamine of *S. cerevisiae* carrying *P. falciparum* DHFR alleles<sup>b</sup> [9] | 3 | $2^3 = 8$ |
| PYR IC50 of *P. vivax* DHFR alleles assayed in *S. cerevisiae*<sup>b</sup> [30] | 4 | $2^4 = 16$ |
| MIC of TEM β-lactamase mutants against 15 β-lactams<sup>b</sup> [43] | 4 | $2^4 = 16^c$ |
| *E. coli* operator binding affinities for 400 repressor sequences [52] | 4 | $4^4 = 256$ |
| Relative fitness among LTEE *E. coli* mutants in DM25<sup>b</sup> [32] | 5 | $2^5 = 32$ |
| Relative fitness among LTEE *E. coli* mutants in DM25 + guanazole<sup>b</sup> [23] | 5 | $2^5 = 32$ |
| HIV replicative capacity on CXCR5+ cells<sup>b</sup> [14] | 5 | $2^5 = 32$ |
| MIC of TEM-β-lactamase mutants against Piperacillin supplemented with clavulanic acid<sup>b</sup> [70] | 6 | $2^5 = 32$ |
| Diploid *S. cerevisiae* mutant growth rate<sup>b</sup> [25] | 6 | $2^6 = 64$ |
| Percent production of minor products by sesquiterpene synthase mutants<sup>b</sup> [47] | 6 | $2^6 = 64$ |
| Percent production of 4-EE by sesquiterpene synthase mutants<sup>b</sup> [47] | 6 | $2^6 = 64$ |
| Percent production of PSD by sesquiterpene synthase mutants<sup>b</sup> [47] | 6 | $2^6 = 64$ |
| 104 mouse DNA-binding proteins' affinity for 10 bp binding motifs [3] | 10 | $4^{10} = 1,048,576^d$ |
| GFP affinity for 10 nucleotide base pair binding motifs [56] | 10 | $4^{10} = 1,048,576$ |
| Affinity of 1,032 DNA-binding proteins spanning eukaryotic diversity against 10 nucleotide base pair binding motifs [76] | 10 | $4^{10} = 1,048,576^e$ |

<sup>a</sup> Written as the product of cardinalities across loci.<sup>b</sup> Another phenotype from this system is included in Table 1.
<sup>c</sup> In total, $16 \times 15$ distinct $\beta$ -lactam compounds = 240 observations are reported in this study.
<sup>d</sup> In total $1,048,576 \times 104$ DNA-binding proteins = 109,051,904 observations are reported in this study.
<sup>e</sup> In total $1,048,576 \times 1,032$ DNA-binding proteins = 1,082,130,432 observations are reported in this study.

Following [74], datasets reporting growth rates [4, 11, 14, 16, 23, 25, 77] or drug-resistance phenotypes [9, 37, 42, 43, 49, 73] were log-transformed before analysis. Following [49], negative two was used in place of log-transformed values when growth rate or drug resistance phenotypes of zero were observed. (In all cases, this is roughly one log order smaller than the smallest non-zero log-transformed value.) In cases where only mean and experimental variances (but not individual replicate observations) were provided, log transformations were approximated by Taylor expansions: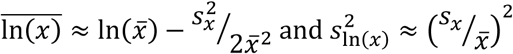. In cases where only means(but not variances) were provided, log transformations were approximated as 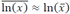.

Following [49], for studies in which experimental variance estimates were provided, we recorded this quantity as a fraction of the total model variance. In one case [9], standard error was reported as standard error over “ at least” two replicates; we therefore assumed *n* = 2 for each observation in that dataset. In one case [32], 95% experimental confidence intervals were reported, so variance estimates were computed under the assumption of normally distributed noise as *s*^2^ = (*n*.CI95/1.96)^2^.

### 2.8 Data and software archiving

Input data files, together with purpose-built MatLab code to perform all analyses described are archived at https://github.com/weinreichlab/JStatPhys2017.Kendall’ s τb correlation coefficient was computed using MatLab code developed elsewhere [10].

## 3 Results

Epistasis can have profound consequences at many levels of biological organization [51, 57, 66, 79]. Here we were particularly interested in the possibility that higher-order epistasis might limit natural selection’ s ability to increase fitness. We indirectly assessed this hypothesis by examining the influence of epistasis on empirical fitness landscape topography as a function of epistatic order (Table 1). As these landscapes are occupied by extant populations that are themselves the product of natural selection, we reasoned that they may be enriched for properties that allow selection to operate efficiently. (See §1.2.)

This study was originally stimulated by Fig. 2 in Palmer et al., 2015 [49], which examined six mutations in the dihydrofolate reductase (DHFR) gene of *E. coli* that contribute to increased resistance to an antimicrobial called pyrimethamine. In that analysis, particular second- and third-order interactions were the third- and second-most influential epistatic terms for fitness landscape topography respectively.Indeed, just two of the first ten most influential epistatic terms were first-order, and in aggregate first-order terms explained just ~28% of the variance in fitness across the landscape. These results seem to challenge the hypothesis outlined in the previous paragraph, and we therefore sought to test its generality using published data from other systems.

Fig. 1 illustrates the application of our analytic pipeline (see Methods) to these same data. Our Fig. 1b closely recapitulates Fig 2a in Palmer et al. 2015 [49]. While the precise sequence of epistatic terms differs slightly (likely because the previous study employed a subtly different framework for computing epistatic terms), higher-order epistatic interactions are again responsible for some the largest reductions in residual variance. Indeed, as previously observed, just two of the first ten terms are first-order, and in aggregate and first-order terms again explain just ~28% of the variance in the data (Table 3a, compare the first two columns with Fig. 2b in [49]).Importantly however, Fig. 1c illustrates that we find a significant, positive correlation between expectation (Eqn. 2) and the observed influence of epistatic terms on landscape topography as a function of their order (τ_b_ = 0.1980, *P* = 0.0377).

**Table 3.** Average epistatic influence on fitness landscape topography as a function of epistatic order in data from [49] and [47]

| Epistatic order | Aggregate reduction in residual variance | Number of epistatic terms significantly different from zero | Mean reduction in residual variance per epistatic term |
| --- | --- | --- | --- |
| **a.** | Log[IC75 of *E. coli* DHFR alleles against trimethoprim] |  |  |
| First | 0.279 | 5 | 0.056 |
| Second | 0.266 | 12 | 0.022 |
| Third | 0.232 | 18 | 0.013 |
| Fourth | 0.144 | 11 | 0.013 |
| Fifth | 0.0684 | 6 | 0.011 |
| Sixth | 0.0065 | 1 | 0.0065 |
| **b.** | Production of 5-epi-aristolochene by sesquiterpene synthase mutants |  |  |
| First | 0.267 | 6 | 0.053 |
| Second | 0.189 | 15 | 0.014 |
| Third | 0.293 | 20 | 0.015 |
| Fourth | 0.118 | 15 | 0.012 |
| Fifth | 0.122 | 6 | 0.024 |
| Sixth | 0.0053 | 1 | 0.0053 |

We next applied our pipeline to 15 other published, combinatorially complete datasets. Results are summarized in Table 1 and shown graphically in Fig. S1. Out of all 16 datasets examined, 14 exhibit a significantly positive correlation between observation and the expectation, and eight of these remain significant after Bonferroni correction for multiple tests. Moreover, across datasets Table 1 exhibits a bias toward small *P* values. Under the null hypothesis (no significant correlation with expectation), we would expect a uniform distribution of *P* values. Instead, the observed distribution is sharply and significantly skewed toward small values (Fig. 2, *G* = 143.77, *P*d.f.=5 ≪ 0.01).

## 4 Discussion

Using a novel analytic pipeline (Fig. 1), we have examined 16 published, combinatorially complete datasets. This analysis broadly confirms our intuition that the influence of epistatic terms on empirical fitness landscape topography should decline with order, i.e., with the number of interacting mutations. While considerable heterogeneity in effect exists among datasets (Table 1), eight of these 16 datasets exhibit a Bonferroni-corrected, significantly positive correlation with expectation (Eqn. 2). And across all 16 datasets, we find a sharp bias toward significant *P* values (Fig. 2). Nor is there any correlation between the size of the dataset and uncorrected *P* value (not shown), suggesting that low statistical power is unlikely to contribute to the overall picture.

The relative magnitudes of epistatic terms depend on the underlying fitness scale employed [33, 74]. Although we log-transforming growth rate and drug resistance data (see §2.7), we have otherwise overlooked this fact. Recently, approaches for systematically rescaling data to minimize higher-order epistatic effects have been introduced [58] (see also [45, 69]). Applications of such methods would certainly have quantitative consequences for results presented here. However, because these approaches (on average) reduce higher-order epistatic terms, we believe this omission renders our conclusions conservative.

We also acknowledge that we failed to honor experimental uncertainty in the sequence of epistatic orders observed, which would almost certainly weaken the signal reported in Table 1. However, our intuition is that this effect would be modest, and moreover, only applies to the nine (of 16) datasets for which experimental uncertainty estimates are available.

### 4.1 The combinatorics of higher-order epistasis

This work was originally stimulated by a previous study [49] that examined six mutations in the DHFR gene responsible for increased pyrimethamine resistance in *E. coli*. Results summarized in Fig. 2 of that study called into question the intuition outlined in §1.2, that higher-order epistasis should only modestly influence naturally occurring fitness landscapes. And the salient features of that figure were recapitulated by our treatment (Fig. 1b, Table 3a).

However, our statistical analysis reveals a strong positive correlation between epistatic influence on fitness topography as a function of epistatic order and that suggested by our evolutionary intuition (Fig. 1c). And applying our analytic pipeline to the aggregated epistatic orders (see §2.6) reported in Fig. 2a of the previous study [49], we again find a significant positive correlation between observed orders and expectation derived from the intuition outlined above (tb = 0.1056, uncorrected *P* = 0.0107). Thus, in this system the substantial influence of some high-order epistatic terms not inconsistent with the idea that high-order epistatic terms should in general only modestly contribute to fitness topography.

The resolution to this puzzle resides in the combinatoric number of epistatic terms. As noted above, given *L* biallelic loci there are 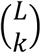 epistatic coefficients of order *k*, and this quantity grows almost exponentially for *k* ≪*L*. Indeed, after normalizing the summed influence of all epistatic terms of order *k* by the number of such terms, we observe that the per-term effect declines almost monotonically in this dataset (Table 3a; see also [74]). This is both as expected on the basis of the evolutionary intuition outlined above and consistent with the statistical analysis of the data in Fig. 1c. A similar picture emerges in our analysis of 5-epi-aristolochene production by sesquiterpene synthase mutants [47]. A number high-order epistatic terms again explain substantial amounts of variance (Fig. S1m), despite a modestly positive correlation coefficient (~0.2) with Eqn. 2 (Table 1). Here again, the resolution reflects the combinatorics of epistatic terms, and the mean per-order effect also declines (Table 3b).

This line of thinking is closely related to the Fourier spectrum of a fitness landscape [45, 63], namely the sum of squared epistatic coefficients as a function of interaction order. (This connection derives formally from Appendix 1, which implies that the squared magnitude of each epistatic coefficient is monotonic in its influence on landscape topography.) The Fourier spectrum is proportional to the binomial coefficient when each genotype’ s fitness is identically and independently distributed. This follows from the fact that on such landscapes all epistatic coefficients are also i.i.d., together with the combinatorics outlined in the previous paragraph. But as already anticipated by results in Table 3, Fourier spectra for both the DHFR and sequiterpine synthase datasets are sharply shifted toward lower-order terms (not shown), as has previously been reported for both sesquiterpene synthase and several others biological datasets [45].

Nevertheless, declining average epistatic effects notwithstanding we find many examples of specific epistatic terms with anomalously large explanatory effects in many of the datasets examined here (Fig. S1). We suggest that these may reflect important mechanistic interactions among those particular mutations in the underlying biology of the system, thus representing potentially fruitful entry points for the molecular biologist [17].

### 4.2 An anthropic perspective and the limits to inferring the effects of natural selection

This study began from the intuition that epistatic influence on empirical fitness landscape topography should decline with epistatic order (see §1.2). This notion rests on three ideas. First, epistasis in general constrains natural selection’ s efficiency [75, 79], and we further speculated that higher-order epistasis might be particularly influential in this respect (see also [59]). Yet datasets such as those in Tables 1 and 2 represent surveys in the genetically local vicinity of extant biological populations. And since biological populations are the product of natural selection, we supposed that they can only persist and succeed if the properties of the fitness landscape enable the efficient action of natural selection [19]. Hence our intuition reflects a line of reasoning that is analogous to the anthropic principle. The anthropic principle holds that the properties of our universe (e.g., the value of its physical constants) should not be regarded as random draws from the space of all conceivable universes. On the contrary, the fact of our observation of those properties sharply constrains our universe to be drawn from the subset of possibilities that are capable of supporting sentient perception.

Does this interpretation of the present analysis imply that natural selection is responsible for the apparently moderate influence of epistatic effects observed? In principle, we might imagine that the strength of epistasis will vary across the exponentially large reaches of sequence space. And if this were true, we might further expect that natural selection would favor populations that find their way to low-epistasis regions on the landscape in preference to those evolving in high-epistasis regions. This would follow if locally reduced epistatic effects sufficiently improved the efficiency of natural selection to allow such populations to outcompete others evolving in regions of the fitness landscape lacking these favorable features. Such population-level competition is sometimes called lineage selection in population genetics [1].

Recently, some support for these ideas has emerged [15, 69]. This work begins from the premise that the epistatic structure of a locally sampled fitness landscapes reflects the way that the constituent mutations were selected by investigators. For example, mutations jointly selected for their ability to confer large fitness gains exhibit more modest epistatic interactions than do mutations whose joint effect is unknown [15]. This finding is at least consistent with lineage selection for reduced epistatic effects.

Critically however, we remain broadly ignorant about the levels of epistasis that exist at random locations in sequence space. Although ever-larger local surveys in sequence space are now becoming possible (e.g., [7, 24, 60]), these are still limited to sparse samples with radii (*L*) of just tens of point mutations. Moreover, the very fact of low (average) high-order epistatic components observed here seems to imply some autocorrelation in epistatic effects across sequence space. This suggests that much wider surveys will be required to develop a sense of what epistasis looks like “ on average.” We thus conclude that for the time being we have only very limited insight into the influence that natural selection has had on the signals detected here.

Indeed, the ability to compare the genetics of what is possible with the genetics of what is observed in nature is generically essential to any demonstration of natural selection [62]. For example, the technological advances described in the previous paragraph are beginning to provide direct experimental access to what is possible at the scale of a handful of mutations, rapidly advancing our ability to detect natural selection acting on first-order mutational effects within individual proteins [7, 21, 62, 67, Wylie et al., in prep]. We look forward to being able to make analogous inferences regarding natural selection’ s influence on epistatic effects in natural populations. (Though these sorts of questions can already be explored using toy fitness landscapes, e.g., [18, 19, 35, 78].)

### 4.3 Epistasis and the efficiency of natural selection

Throughout, we have assumed that high-order epistatic interactions reduce the efficiency of natural selection (§1.2). Our observation that the influence of epistatic terms on naturally occurring fitness landscapes declines with epistatic order represents an indirect test of this idea. However, we lack a detailed theoretical understanding of this connection.

Perhaps the most well-developed results concern the influence of epistasis on the selective accessibility of mutational trajectories to high fitness genotypes. First, sign epistasis means that the sign of the fitness of a mutation of selection varies with genetic background [75], and it renders selectively inaccessible at least some mutational trajectories to high fitness (e.g., [73]). But connections between sign epistasis and epistatic order are only now being developed [13]. Second, a subsetting approach similar to ours (§2.3) was recently used to examine the influence of epistatic interactions selectively accessible mutational trajectories to high fitness genotypes [59] in six of the datasets described here. Those authors found that higher-order terms indeed substantially alter the identity of selectively favored mutational trajectories to high-fitness genotypes, as well as their probabilities of realization. Further and consistent with findings here, that study also noted that the absolute magnitude of epistatic terms had an even larger effect on realized mutational trajectories than did their interaction order.

However, epistasis has long been understood to influence not just the selectively accessibility of high fitness genotypes but also the pace at which natural selection both increases the frequency of beneficial mutations (e.g., [20]) and at which it purges deleterious mutations (e.g., [34]). This work is closely related to the role that genetic recombination can play in “ unlocking” epistatically interacting mutations (e.g., [5, 44]). To our knowledge the relationship between these effects and higher-order epistasis remains entirely unexplored.

In addition, we have only quantitatively examined the sequence of epistatic orders sorted by explanatory power (Fig. 1c). Thus, a great deal of information present in these data (e.g., the slopes in Figs. 1b and S1) remains to be examined. And of course, the number and size of available combinatorially complete datasets continues to grow, motivating further work in this regard. It seems reasonable to suppose that the development and testing of more nuanced theoretical predictions may be possible using data of the sort examined here.

Finally, we note that the Fourier-Walsh framework employed here depends on the availability of combinatorially complete datasets. But the experimental demands of this approach grow exponentially with the number of mutations examined. This fact sharply limits the scalability of analytic pipelines like ours. Recently, theoretical progress has been made in the analysis of less-than-complete datasets [6, 13], and older work has also explored this idea [27, 64]. Theory that allows inferences using sparse datasets is likely to be a key advance in our ability explore broad, evolutionarily fascinating questions such as those considered here.

## Acknowledgements

We are grateful to Tony Dean, David Hall, Sebastian Matuszewski, and Vaughn Cooper for providing raw data files. We also acknowledge constructive feedback on an earlier draft of this manuscript from Guillaume Achaz, Kristina Crona, Inês Fragata, Joachim Krug, Sebastian Matuszewski and Brandon Ogbunugafor. DMW is supported in part by National Science Foundation Grant DEB-1556300 and National institutes of Health Grant R01GM095728. RBH is supported in part by the National Science Foundation under Cooperative Agreement No. DBI-0939454.

## Appendix 1 The explanatory power of Fourier-Walsh coefficients is monotonic in their absolute magnitude

Assume two fitness functions defined over *L* biallelic loci are represented as column vectors 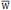 and 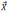with Fourier-Walsh coefficients 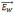 and 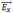. Define the sum of squares distance between 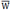 and 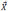 as 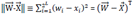. 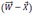 where *w*_*i*_ and *x*_*i*_are the *i*^th^ components of 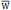and 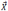 respectively.

### Theorem 1: Sum of squares distance equivalence

Proof: By definition

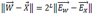

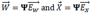

where **Ψ** is the Hadamard matrix (see §2.2). Therefore

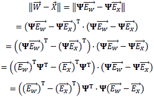

But recall that **Ψ ^T^Ψ** = 2^L^**I**, where **I** is the identity matrix. Thus

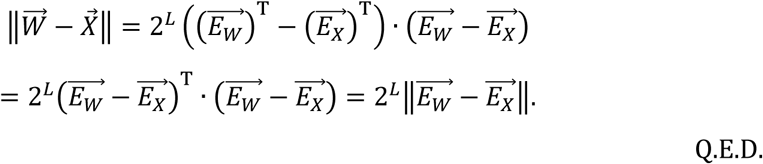

An interesting property of the Hadamard matrix is that **Ψ**^T^ = 2^*L*^ **Ψ**^-1^. Without the 2^*L*^ this equality is the hallmark of a rotational transformation. This means that Fourier-Walsh coefficients are simply the result of a high dimensional axis rotation of the coordinates of function space, together with a uniform contraction. This provides intuition into Theorem 1: rotating the space and contracting it uniformly only changes the distance between two vectors in the space by the constant of contraction.

### Theorem 2: Minimizing the sum-of-squares distance of subsetting approximations

The subsetting approximation 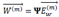 that minimizes the sum of squares distance to function 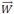is the one whose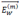 uses the *m* largest components in absolute value in 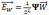.

**Proof:** By Theorem 1, the sum of squares distance between 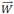 and 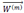 is 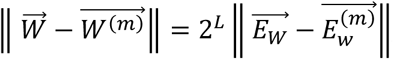, which means that we can equivalently solve the minimization problem on either side of the equality. And trivially, the right-hand side is minimized when the *m* nonzero components in 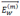 are the *m* largest components in absolute value in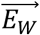 (The squaring of differences in epistatic terms in the definition of 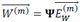 removes the significance of their sign.)

**Fig. S1.**
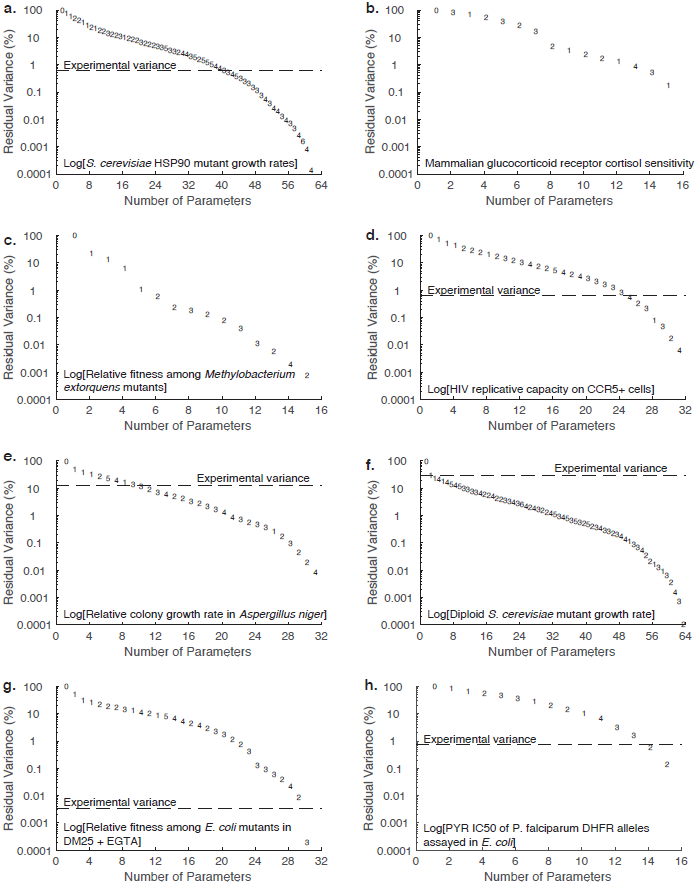
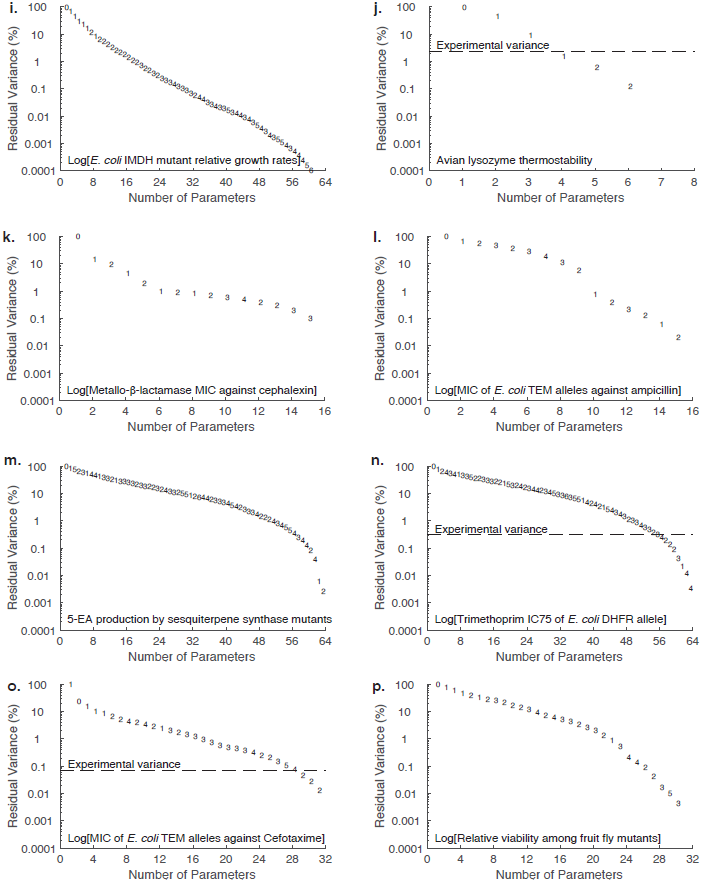
Predicted fitness under a succession of models using only the *m* largest epistatic terms. As in Fig. 1b, the symbols represent the epistatic order of each successive parameter added to the model, and experimental variance (where provided) expressed as a fraction of total variance. Citations: **a.** [4], **b.** [8], **c.** [11], **d.** [14], **e.** [16], **f.** [25], **g.** [23], **h.** [37], **i.** [38], **j.** [39], **k.** [42], **l.** [43], **m.** [47], **n.** [49], see also Fig. 1, **o.** [73]**, p.** [77].

## References

1. Akçay, E. and J. Van Cleve: There is no fitness but fitness, and the lineage is its bearer. Philosophical Transactions of the Royal Society B: Biological Sciences 371, (2016)

2. Avery, L. and S. Wasserman: Ordering gene function: the interpretation of epistasis in regulatory hierarchies. Trends in Genetics 8, 312–316 (1992)

3. Badis, G., M.F. Berger, A.A. Philippakis, S. Talukder, A.R. Gehrke, S.A. Jaeger, E.T. Chan, G. Metzler, A. Vedenko, X. Chen, H. Kuznetsov, C.-F. Wang, D. Coburn, D.E. Newburger, Q. Morris, T.R. Hughes, and M.L. Bulyk: Diversity and Complexity in DNA Recognition by Transcription Factors. Science (2009)

4. Bank, C., S. Matuszewski, R.T. Hietpas, and J.D. Jensen: On the (un)predictability of a large intragenic fitness landscape. Proceedings of the National Academy of Sciences 113, 14085–14090 (2016)

5. Barton, N.H.: Why Sex and Recombination? Cold Spring Harbor Symposia on Quantitative Biology 74, 187–195 (2009)

6. Beerenwinkel, N., L. Lpachter, and B. Sturmfels: Epistasis and shapes of fitness landscapes. Statistica Sinica 17, 1317–1342 (2007)

7. Boucher, J.I., P. Cote, J. Flynn, L. Jiang, A. Laban, P. Mishra, B.P. Roscoe, and D.N.A. Bolon: Viewing Protein Fitness Landscapes Through a Next-Gen Lens. Genetics 198, 461–471 (2014)

8. Bridgham, J.T., E.A. Ortlund, and J.W. Thornton: An epistatic ratchet constrains the direction of glucocorticoid receptor evolution. Nature 461, 515–519 (2009)

9. Brown, K.M., M.S. Costanzo, W. Xu, S. Roy, E.R. Lozovsky, and D.L. Hartl: Compensatory mutations restore fitness during the evolution of dihydrofolate reductase. Molecular Biology and Evolution 27, 2682–2690 (2010)

10. Burkey, J. A non-parametric monotonic trend test computing Mann-Kendall Tau, Tau-b, and Sen’ s Slope written in Mathworks-MATLAB implemented using matrix rotations. 2006.

11. Chou, H.-H., H.-C. Chiu, N.F. Delaney, D. Segrè, and C.J. Marx: Diminishing returns epistasis among beneficial mutations decelarates adaptation. Science 322, 1190–1192 (2011)

12. Costanzo, M.S., K.M. Brown, and D.L. Hartl: Fitness trade-offs in the evolution of dihydrofolate reductase and drug rsistance in Plasmodium falciparum. PLoS One 6, e19636 (2011)

13. Crona, K., A. Gavryushkin, D. Greene, and N. Beerenwinkel: Inferring Genetic Interactions From Comparative Fitness Data. bioRxiv (2017)

14. da Silva, J., M. Coetzer, R. Nedellec, C. Pastore, and D.E. Mosier: Fitness Epistasis and Constraints on Adaptation in a Human Immunodeficiency Virus Type 1 Protein Region. Genetics 185, 293–303 (2010)

15. de Visser, J.A.G.M. and J. Krug: Empirical fitness landscapes and the predictability of evolution. Nat Rev Genet 15, 480–490 (2014)

16. de Visser, J.A.G.M., S.-C. Park, and J. Krug: Exploring the Effect of Sex on Empirical Fitness Landscapes. American Naturalist 174, S15–S30 (2009)

17. Dean, A.M. and J.W. Thornton: Mechanistic approaches to the study of evolution: the functional synthesis. Nature Reviews genetics 8, 675–688 (2007)

18. Draghi, J.A., T.L. Parsons, G.P. Wagner, and J.B. Plotkin: Mutational robustness can facilitate adaptation. Nature 463, 353–355 (2010)

19. Draghi, J.A. and J.B. Plotkin: Selection biases the prevalence and type of epistasis along adaptive trajectories. Evolution 67, 3120–3131 (2013)

20. Eshel, I. and M.W. Feldman: On the evolutionary effect of recombination. Theoretical Population Biology 1, 88–100 (1970)

21. Firnberg, E., J.W. Labonte, J.J. Gray, and M. Ostermeier: A Comprehensive, High-Resolution Map of a Gene’ s Fitness Landscape. Molecular Biology and Evolution 31, 1581–1592 (2014)

22. Fisher, R.A.: The genetical theory of natural selection. Clarendon Press, Oxford, UK (1930)

23. Flynn, K.M., T.F. Cooper, F.B.G. Moore, and V.S. Cooper: The Environment Affects Epistatic Interactions to Alter the Topology of an Empirical Fitness Landscape. PLOS Genetics 9, e1003426 (2013)

24. Fowler, D.M. and S. Fields: Deep mutational scanning: a new style of protein science. Nat Meth 11, 801–807 (2014)

25. Hall, D.W., M. Agan, and S.C. Pope: Fitness epistasis among 6 biosynthtic loci in the budding yeast Saccharomyces cervisiae. Journal of Heredity 1010, S75–S84 (2010)

26. Heckendorn, R.B. and D. Whitley: Predicting epistasis from mathematical models. Evolutionary Computation 7, 69–101 (1997)

27. Heckendorn, R.B. and A.H. Wright: Efficient linkage discovery by limited probing. Evolutionary Computation 12, 517–545 (2004)

28. Holm, S.: A simple sequentially rejective multiple test procedure. Scandinavian Journal of Statistics 6, 65–70 (1979)

29. Iwasa, Y., F. Michor, and M.A. Nowak: Stochastic tunnels in evolutionary dynamics. Genetics 166, 1571–1579 (2004)

30. Jiang, P.-P., R.B. Corbett-Detig, D.L. Hartl, and E.R. Lozovsky: Accessible Mutational Trajectories for the Evolution of Pyrimethamine Resistance in the Malaria Parasite Plasmodium vivax. Journal of Molecular Evolution 77, 81–91 (2013)

31. Kendall, M.G.: A new measure of rank correlation. Biometrika 30, 81–93 (1938)

32. Khan, A.I., D.M. Dinh, D. Schneider, R.E. Lenski, and T.F. Cooper: Negative epistasis between beneficial mutations in an evolving bacterial population. Science 332, 1193–1196 (2011)

33. Knies, J.L., F. Cai, and D.M. Weinreich: Enzyme Efficiency but Not Thermostability Drives Cefotaxime Resistance Evolution in TEM-1 β-Lactamase. Molecular Biology and Evolution 34, 1040–1054 (2017)

34. Kondrashov, A.S.: Deleterious mutations and the evolution of sex. Nature 336, 435–440 (1988)

35. Lan, Y., A. Trout, D.M. Weinreich, and C.S. Wylie: Natural selection can favor the evolution of ratchet robustness over evolution of mutational robustness. bioRxiv (2017)

36. Leem, S., H.-h. Jeong, J. Lee, K. Wee, and K.-A. Sohn: Fast detection of high-order epistatic interactions in genome-wide association studies using information theoretic measure. Computational Biology and Chemistry 50, 19–28 (2014)

37. Lozovsky, E.R., T. Chookajorn, K.M. Brown, M. Imwong, P.J. Shaw, S. Kamchonwongpaisan, D.E. Neafsey, D.M. Weinreich, and D.L. Hartl: Stepwise acquisition of pyrimethamine resistance in the malaria parasite. Proceedings of the National Academy of Sciences 106, 12025–12030 (2009)

38. Lunzer, M., S.P. Miller, R. Felsheim, and A.M. Dean: The biochemical architecture of an ancient adaptive landscape. Science 310, 499–501 (2005)

39. Malcolm, B.A., K.P. Wilson, B.W. Matthews, J.F. Kirsch, and A.C. Wilson: Ancestral lysozymes reconstructed, neutrality tested, and thermostability linked to hydrocarbon packing. Nature 345, 86–89 (1990)

40. Maynard Smith, J.: Natural selection and the concept of a protein space. Nature 225, 563–565 (1970)

41. McCandlish, D.M.: Visualizing fitness landscapes. Evolution 65, 1544–1558 (2011)

42. Meini, M.-R., P.E. Tomatis, D.M. Weinreich, and A.J. Vila: Quantitative Description of a Protein Fitness Landscape Based on Molecular Features. Molecular Biology and Evolution 32, 1774–1787 (2015)

43. Mira, P.M., J.C. Meza, A. Nandipati, and M. Barlow: Adaptive Landscapes of Resistance Genes Change as Antibiotic Concentrations Change. Molecular Biology and Evolution (2015)

44. Neher, R.A. and B.I. Shraiman: Competition between recombination and epistasis can cause a transition from allele to genotype selection. Proceedings of the National Academy of Sciences 106, 6866–6871 (2009)

45. Neidhart, J., I.G. Szendro, and J. Krug: Exact results for amplitude spectra of fitness landscapes. Journal of Theoretical Biology 332, 218–227 (2013)

46. Nelson, R.M., M. Kierczak, and Ö. Carlborg: Higher Order Interactions: Detection of Epistasis Using Machine Learning and Evolutionary Computation. In: C. Gondro, J. van der Werf, and B. Hayes(eds.) Genome-Wide Association Studies and Genomic Prediction, pp. 499–518. Humana Press, Totowa, NJ (2013)

47. O’Maille, P.E., A. Malone, N. Dellas, B.A. Hess, Jr., L. Smentek, I. Sheehan, B.T. Greenhagen, J. Chappell, G. Manning, and J.P. Noel: Quantitative exploration of the catalytic landscape separating divergent plant sesquiterpene synthases. Nature Chemical Biology 4, 617–623 (2008)

48. Orr, H.A.: Fitness and its role in evolutionary genetics. Nat Rev Genet 10, 531–539 (2009)

49. Palmer, A.C., E. Toprak, M. Baym, S. Kim, A. Veres, S. Bershtein, and R. Kishony: Delayed commitment to evolutionary fate in antibiotic resistance fitness landscapes. Nature Communications 6, 7385 (2015)

50. Phillips, P.C.: The language of gene interaction. Genetics 149, 1167–1171 (1998)

51. Phillips, P.C.: Epistasis — the essential role of gene interactions in the structure and evolution of genetic systems. Nature Reviews genetics 9, 855–867 (2008)

52. Poelwijk, F., D.J. Kiviet, and S.J. Tans: Evolutionary potential of a duplicated repressor-operator pair: Simulating pathways using mutational data. PLoS Computational Biology 2, e58 (2006)

53. Poelwijk, F., D.J. Kiviet, D.M. Weinreich, and S.J. Tans: Empirical fitness landscapes reveal accessible evolutionary paths. Nature 445, 383–386 (2007)

54. Poelwijk, F.J., V. Krishna, and R. Ranganathan: The Context-Dependence of Mutations: A Linkage of Formalisms. PLoS Comput Biol 12, e1004771 (2016)

55. Provine, W.B.: Sewall Wright and Evolutionary Biology. University of Chicago Press, Chicago (1986)

56. Rowe, W., M. Platt, D.C. Wedge, P.J. Day, D.B. Kell, and J. Knowles: Analysis of a complete DNA–protein affinity landscape. Journal of The Royal Society Interface 7, 397–408 (2010)

57. Sackton, T.B. and D.L. Hartl: Genotypic Context and Epistasis in Individuals and Populations. Cell 166, 279–287 (2016)

58. Sailer, Z.R. and M.J. Harms: Detecting High-Order Epistasis in Nonlinear Genotype-Phenotype Maps. Genetics 205, 1079–1088 (2017)

59. Sailer, Z.R. and M.J. Harms: High-order epistasis shapes evolutionary trajectories. PLOS Computational Biology 13, e1005541 (2017)

60. Sarkisyan, K.S., D.A. Bolotin, M.V. Meer, D.R. Usmanova, A.S. Mishin, G.V. Sharonov, D.N. Ivankov, N.G. Bozhanova, M.S. Baranov, O. Soylemez, N.S. Bogatyreva, P.K. Vlasov, E.S. Egorov, M.D. Logacheva, A.S. Kondrashov, D.M. Chudakov, E.V. Putintseva, I.Z. Mamedov, D.S. Tawfik, K.A. Lukyanov, and F.A. Kondrashov: Local fitness landscape of the green fluorescent protein. Nature 533, 397–401 (2016)

61. Sokal, R.R. and F.J. Rohlf: Biometry. W.H. Freeman and Company, New York (1995)

62. Springer, S.A., M. Manhart, and A.V. Morozov: Separating Spandrels from Phenotypic Targets of Selection in Adaptive Molecular Evolution. In: P. Pontarotti(eds.) Evolutionary Biology: Convergent Evolution, Evolution of Complex Traits, Concepts and Methods, pp. 309–325. Springer International Publishing, Cham (2016)

63. Stadler, P.F.: Landscapes and their correlation functions. Journal of Mathematical Chemistry 20, 1–45 (1996)

64. Stadler, P.F.: Spectral landscape theory. In: J.P. Crutchfield and P. Schuster(eds.) Evolutionary dynamics: Exploring the interplay of selection, accident, neutrality, and function, pp. 221–272. Oxford University Press, Oxford (2003)

65. Stadler, P.F. and R. Happel: Random field models for fitness landscapes. Journal of Mathematical Biology 38, 435–478 (1999)

66. Starr, T.N. and J.W. Thornton: Epistasis in protein evolution. Protein Science 25, 1204–1218 (2016)

67. Stiffler, Michael A., Doeke R. Hekstra, and R. Ranganathan: Evolvability as a Function of Purifying Selection in TEM-1 β-Lactamase. Cell 160, 882–892 (2015)

68. Sun, X., Q. Lu, S. Mukherjee, P.K. Crane, R. Elston, and M.D. Ritchie: Analysis pipeline for the epistasis search – statistical versus biological filtering. Frontiers in Genetics 5, (2014)

69. Szendro, I.G., M. Schenk, J. Franke, J. Krug, and J.A.G.M. de Visser: Quantitative analyses of empirical fitness landscapes. Journal of Statistical Mechanics P01, 005 (2013)

70. Tan, L., S. Serene, H.X. Chao, and J. Gore: Hidden randomness between fitness landscapes limits reverse evolution. Physical Review Letters 106, 198102 (2011)

71. Weinberger, E.D.: Fourier and Taylor series on fitness landescapes. Biological Cybernetics 65, 321–330 (1991)

72. Weinreich, D.M. and L. Chao: Rapid evolutionary escape by large populations from local fitness peaks is likely in nature. Evolution 59, 1175–1182 (2005)

73. Weinreich, D.M., N.F. Delaney, M.A. DePristo, and D.L. Hartl: Darwinian evolution can follow only very few mutational paths to fitter proteins. Science 312, 111–114 (2006)

74. Weinreich, D.M., Y. Lan, C.S. Wylie, and R.B. Heckendorn: Should Evolutionary Geneticists Worry about High Order Epistasis? Current Opinion in Development and Genetics 23, 700–707 (2013)

75. Weinreich, D.M., R.A. Watson, and L. Chao: Perspective: Sign epistasis and genetic constraint on evolutionary trajectories. Evolution 59, 1165–1174 (2005)

76. Weirauch, Matthew T., A. Yang, M. Albu, A.G. Cote, A. Montenegro-Montero, P. Drewe, Hamed S. Najafabadi, Samuel A. Lambert, I. Mann, K. Cook, H. Zheng, A. Goity, H. van Bakel, J.-C. Lozano, M. Galli, M.G. Lewsey, E. Huang, T. Mukherjee, X. Chen, John S. Reece-Hoyes, S. Govindarajan, G. Shaulsky, Albertha J.M. Walhout, F.-Y. Bouget, G. Ratsch, Luis F. Larrondo, Joseph R. Ecker, and Timothy R. Hughes: Determination and Inference of Eukaryotic Transcription Factor Sequence Specificity. Cell 158, 1431–1443

77. Whitlock, M.C. and D. Bourguet: Factors affecting the genetic load in Drosophila: Synergistic epistasis and correlations among fitness components. Evolution 54, 1654–1660 (2000)

78. Wilke, C.O.: Selection for fitness versus selection for robustness in RNA secondary structure folding. Evolution 55, 2412–2420 (2001)

79. Wolf, J.B., E.D.I. Brodie, and M.J. Wade, eds. Epistasis and the Evolutionary Process. 2000, Oxford University Press: New York. 330.

80. Wright, S.: Evolution in Mendelian Populations. Genetics 16, 97–159 (1931)

81. Wright, S.: The roles of mutation, inbreeding, crossbreeding and selection in evolution. In: D.F. Jones(eds.) Proceedings of the Sixth International Congress of Genetics, pp. 356–366. Brooklyn Botanic Garden, Menasha, WI (1932)

